# Comparative genomics of the sexually transmitted parasite *Trichomonas vaginalis* reveals relaxed and convergent evolution and genes involved in spillover from birds to humans

**DOI:** 10.1101/2024.12.22.629724

**Authors:** Steven A. Sullivan, Jordan C. Orosco, Francisco Callejas-Hernández, Frances Blow, Hayan Lee, Timothy Ranallo-Benavidez, Andrew Peters, Shane Raidal, Yvette A. Girard, Christine K. Johnson, Krysta Rogers, Richard Gerhold, Hayley Mangelson, Ivan Liachko, Harsh Srivastava, Chris Chandler, Daniel Berenberg, Richard A. Bonneau, Po-Jung Huang, Yuan-Ming Yeh, Chi-Ching Lee, Hsuan Liu, Petrus Tang, Ting-Wen Chen, Michael C. Schatz, Jane M. Carlton

## Abstract

*Trichomonas vaginalis* is the causative agent of the venereal disease trichomoniasis which infects men and women globally and is associated with serious outcomes during pregnancy and cancers of the human reproductive tract. Trichomonads parasitize a range of hosts in addition to humans including birds, livestock, and domesticated animals. Recent genetic analysis of trichomonads recovered from columbid birds has provided evidence that these parasite species undergo frequent host-switching, and that a current epoch spillover event from columbids likely gave rise to *T. vaginalis* in humans. We undertook a comparative evolutionary genomics study of seven trichomonad species, generating chromosome-scale reference genomes for *T. vaginalis* and its avian sister species *Trichomonas stableri*, and assemblies of five other species that infect birds and mammals. Human-infecting trichomonad lineages have undergone recent and convergent genome size expansions compared to their avian sister species, and the major contributor to their increased genome size is increased repeat expansions, especially multicopy gene families and transposable elements, with genetic drift likely a driver due to relaxed selection. Trichomonads have independently host-switched twice from birds to humans, and genes implicated in the transition to the human host include those associated with host tissue adherence and phagocytosis, extracellular vesicles, and CAZyme virulence factors.

## Introduction

Trichomoniasis, the most prevalent non-viral venereal disease of humans, is caused by the protozoan *Trichomonas vaginalis* that infects the lower genital tract of men (urethra and prostate) and women (vulva, vagina, and cervix). Symptoms include foul-smelling vaginal discharge and genital itching, and infections are associated with an increased risk of cervical and prostate cancer, HIV infection, and complications during pregnancy^1^. Other human-infecting trichomonads include the oral parasite *Trichomonas tenax* associated with periodontal disease^2^, and the intestinal parasite *Pentatrichomonas hominis* associated with gastrointestinal distress and diarrhea^3^. Trichomonad species also infect a wide range of vertebrate hosts, including birds, livestock, and domesticated animals. Avian trichomonads include *Trichomonas gallinae*^4^ which infects the upper gastrointestinal (GI) tract of a diversity of birds including doves, pigeons, and songbirds as well as raptors that may prey on infected birds. *Trichomonas* species have also been documented as an important cause of morbidity and mortality in Passeriformes (perching birds) and responsible for a large decline of greenfinch and chaffinch populations in Great Britain at the beginning of the 21^st^ century^5^. In 2008, novel parasites with genetic markers highly similar to *T. vaginalis* (dubbed *“T. vaginalis*-like*”*) were reported in white-winged doves and mourning doves from Arizona and Texas, and in Pacific coast band-tailed pigeons from California^6^. *T. vaginalis*-like parasites recovered from the latter during a 2011-12 outbreak in California were analyzed in detail and given the species name *Trichomonas stableri*^7^. *T. vaginalis* most likely originated as a zoonosis from American pigeons and doves during a recent spillover event during the current (Holocene) epoch following human colonization of the Americas^8^. It has been hypothesized that its ancestor moved from the upper GI tract of columbids into the human reproductive tract via barrier contraceptives or more commonly through human contact with bird-infected water^8^.

Genome-scale interrogation of *T. vaginalis* biology and evolution began in 2007 with the generation of the first genome assembly of strain G3 using Sanger sequencing^9^. The genome was extremely difficult to assemble and annotate due to its repetitive DNA content comprised of thousands of highly similar transposable elements (TEs) and many multicopy gene families, and its unexpectedly large size (∼180 Mb) compared to genomes of other human parasitic protists^10^. Especially disruptive to genome assembly were thousands of newly discovered Maverick sequences: long (∼10-28 Kb) virus-like DNA transposons found (typically in much lower abundance) in all major eukaryotic lineages except plants and mammals^11^. The resulting *T. vaginalis* G3 draft genome assembly of consisted of >64,000 scaffolds and contigs, and while fragmentation made it inadequate for accurate counting of repetitive elements, the assembly generated new insights into (1) multicopy gene families involved in the parasite’s active endocytic and phagocytic life-style such as protein kinases, peptidases, and membrane trafficking proteins; (2) surface proteins including the saposin-like (SAPLIP) pore-forming cytolytic family and highly diverse BspA-like proteins that may mediate parasite adherence to vaginal epithelial cells required to establish and maintain an infection; and (3) novel metabolic pathways shaped by putative prokaryote-to-eukaryote lateral gene transfer events. With few exceptions, other trichomonad genomes remained unsequenced, although in the interim, molecular phylogenies showed avian trichomonads to be the closest known relatives of *T. vaginalis* and *T. tenax*^6, 12^, with columbids inferred to be the ancestral host of the genus, and the source of at least two independent host switches to humans^8^. Host switching has been posited to be a strong macroevolutionary force in genus *Trichomonas*^8^.

The high copy numbers of *T. vaginalis* gene families that Carlton et al., reported in 2007^9^ raised the question of whether such repeat expansions persist because they are adaptive or because of other evolutionary mechanisms such as drift. Selection for a paralog with a new function (neofunctionalization) or for increased gene dosage can contribute to long-term preservation of gene duplicates^13^. Differential expression of paralogs in different environments, e.g., different host species or tissue, has been cited to infer adaptive new roles for some gene duplicates; such evidence has been reported in *T. vaginalis* for some paralogs in some multicopy gene families, such as cysteine proteases (reviewed in^10^). However, such evidence exists for just a small fraction of paralogs in gene families overall, and adaptive processes such as positive or purifying selection are unlikely to explain the high burden of TEs in *T. vaginalis* -- generally assumed to be deleterious invaders of exogenous origin. Alternatively, when selection is relaxed -- as when functional constraints on a gene are removed, or effective population size is reduced, both of which can occur when an organism switches host environments -- its converse, genetic drift, comes to the fore, randomly fixing or deleting alleles. Elevated levels of drift are predicted by the ‘mutation hazard hypothesis’^14^ to (among other things) increase sequence copy number, including genes and TEs. Such hypotheses have not been tested in trichomonads before.

To expand the number of available trichomonad genome sequences, address key knowledge gaps such as those noted above, and identify genes implicated in the spillover event from avian to human host, we leveraged long-read and chromosomal conformation sequencing to generate chromosome-scale reference genome assemblies for *T. vaginalis* G3 and its sister species the avian parasite *T. stableri*. We used short-read sequencing to also assemble draft genomes of two other human-infecting species, *T. tenax* and *P. hominis,* and three other bird-infecting species. These seven assembled genomes represent the most extensive whole genome sequence dataset of trichomonads to date, and enabled unprecedented comparative genomics, including estimate of gene and TE content in closely-related species from different hosts, and visualization of synteny. We offer new insights into trichomonad evolution, including evidence for relaxed selection accompanying the inferred host switch from birds in two human-infective *Trichomonas* lineages, which likely explains the striking genome size variation among these trichomonads. We additionally identify convergently evolving genes in human-infecting species that were putatively involved in the transition from bird to human host.

## Results

### Comprehensive TE annotation in *T. vaginalis* from a new chromosome-scale reference assembly

We generated a new, chromosome-scale reference assembly of *T. vaginalis* strain G3 using Pacific Bioscience long-read sequencing augmented with chromosome conformation capture (‘PacBio/Hi-C’). The new assembly comprises six chromosome-scale scaffolds matching the published *T. vaginalis* karyotype number^15^ ranging from 20-40 Mb and ∼177 Mb total length (in contrast to the 2007^9^ Sanger assembly of >64,000 scaffolds [range 0.2-585 kb] and 176 Mb total length). Microsatellite and rRNA loci localized to metaphase chromosome squashes by FISH^9, 16^ were mapped to the assembly to assign chromosome numbers I-VI to the scaffolds (**Figure 1**). We improved the accuracy of the *T. vaginalis* predicted proteome, identifying 37,794 protein-coding genes (**Table 1**), with 46% annotated as ‘hypothetical’ (compared to the 59,681 genes with 75% hypotheticals predicted in 2007^9^). Improved annotation of the 16 major multicopy gene families, many of which are associated with cell surface activity, parasite-host interaction, and the degradome, increased their copy numbers, more than quadrupling it in the case of cysteine peptidase Clan CA, family C1 (**Supplementary Table 1**). The >600 rDNA genes identified in 2007 collapsed to eleven 28S/5.8S/18S rRNA cassettes tandemly arrayed on chromosome II, agreeing with FISH results^9^ (**Figure 1**). We extended a previously reported^17^ block of genes laterally transferred from a relative of the firmicute bacterium *Peptoniphilus harei*, from 37 Kb containing 27 genes to 47 Kb containing 45 genes (**Supplementary Table 2**). The *T. vaginalis* genome remains densely packed with protein-coding genes and TEs, with an average length of 1,131 bp (median length 520 bp) between them.

**Figure 1.**
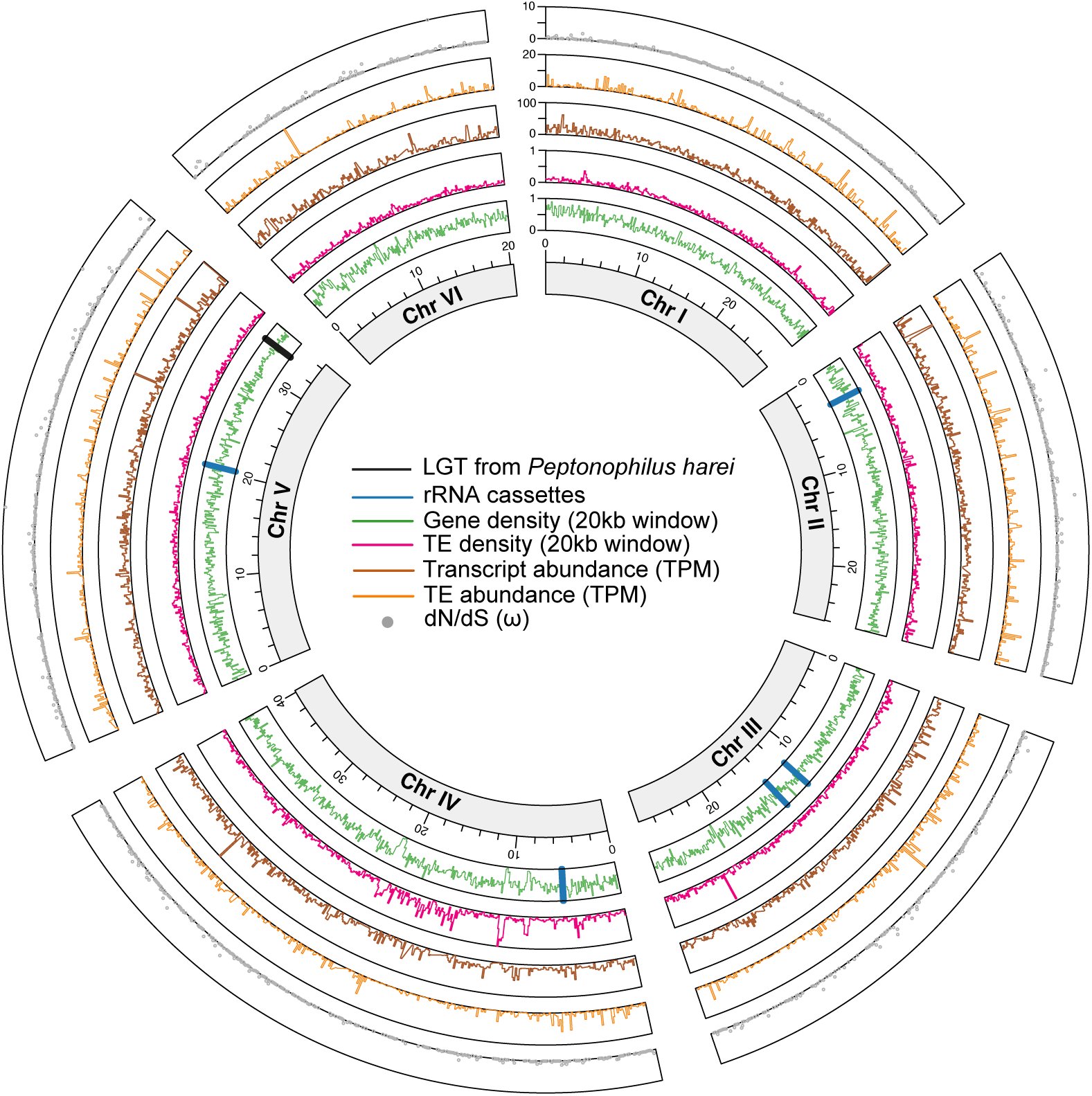
Architecture and genome features of *T. vaginalis* G3 across its six chromosomes. The concentric rings, from innermost to outermost, represent: (1) chromosome size in Mb; (2) gene density (green plot) shown in 20 Kb windows; vertical blue lines represent 11 rRNA cassettes, and the vertical black line represents the 47.5 Kb block from an LGT event of the bacterium *Peptoniphilus harei*; (3) TE density (pink plot) shown in 20 Kb windows; (4) transcript abundance (brown plot) of all genes shown as transcripts per million in 100 Kb windows; (5) TE transcript abundance (orange plot) of annotated TE genes) in 100 Kb windows; and (6) dN/dS values (grey dots). The axes are shown next to chromosome 1.

**Table 1.**
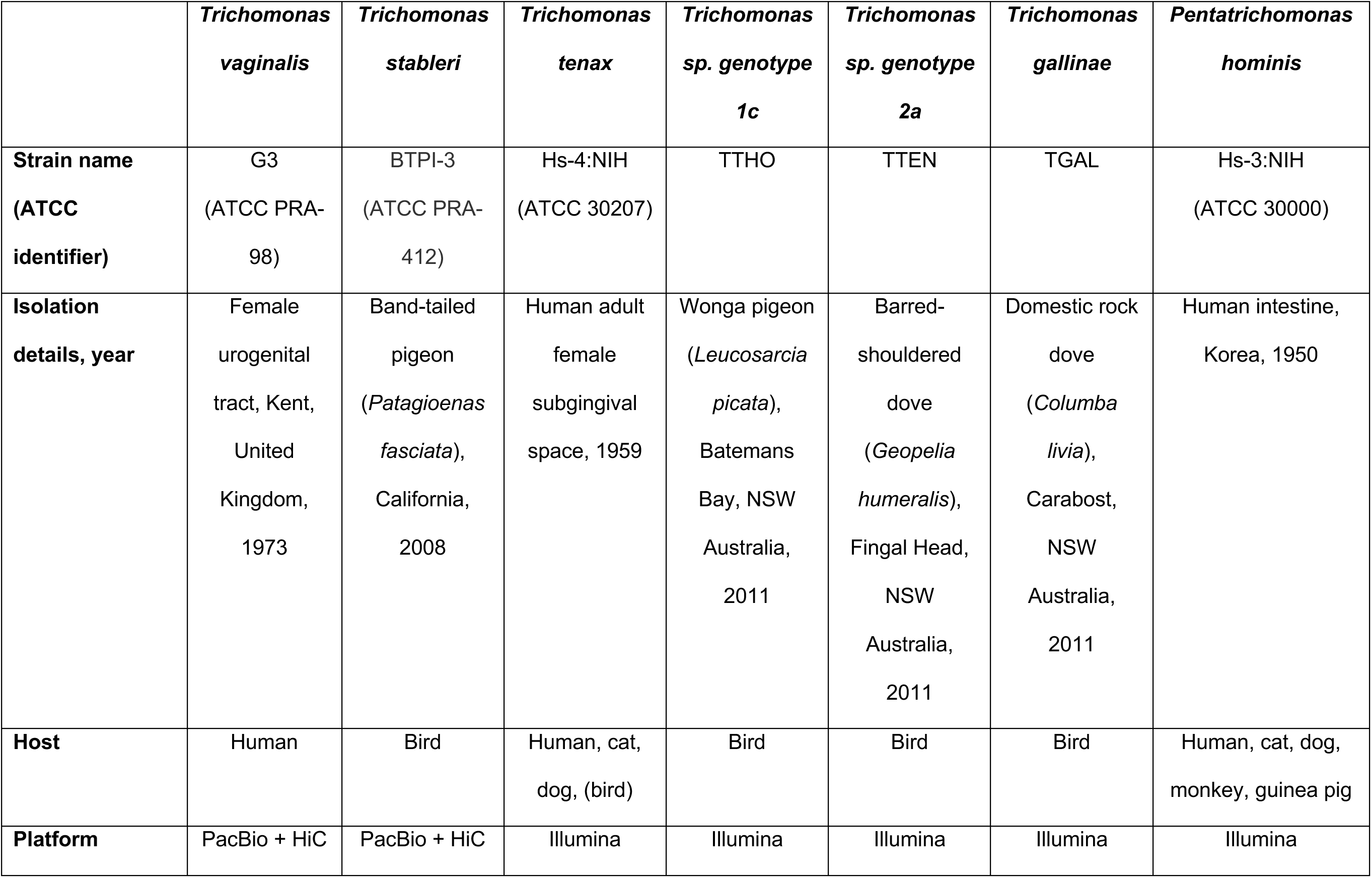
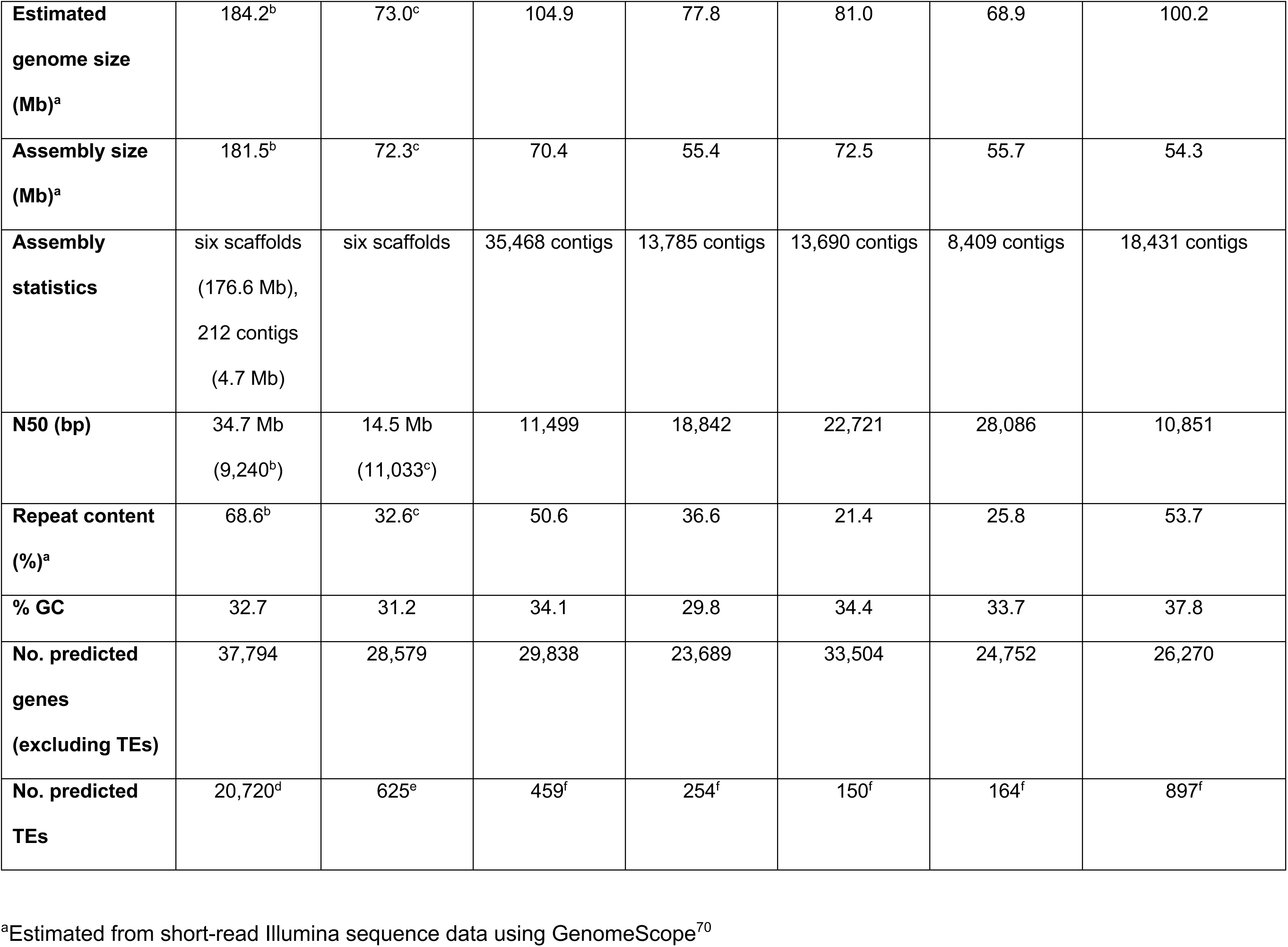
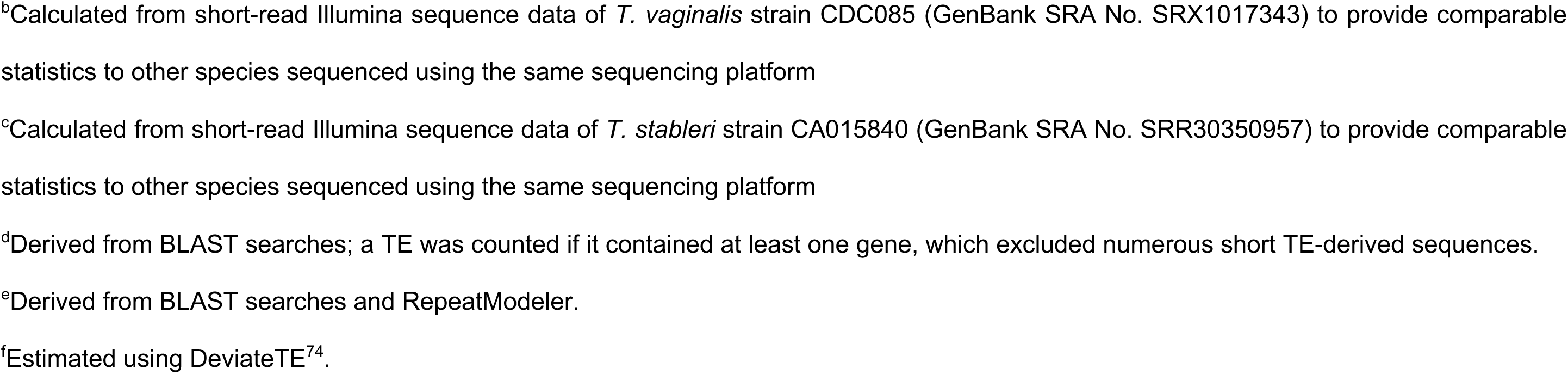
Genome assembly and annotation statistics for seven trichomonad species. ATCC: American Type Culture Collection.

|  | *Trichomonas vaginalis* | *Trichomonas stableri* | *Trichomonas tenax* | *Trichomonas sp. genotype 1c* | *Trichomonas sp. genotype 2a* | *Trichomonas gallinae* | *Pentatrichomonas hominis* |
| --- | --- | --- | --- | --- | --- | --- | --- |
| **Strain name (ATCC identifier)** | G3<br>(ATCC PRA-98) | BTPI-3<br>(ATCC PRA-412) | Hs-4:NIH<br>(ATCC 30207) | TTHO | TTEN | TGAL | Hs-3:NIH<br>(ATCC 30000) |
| **Isolation details, year** | Female urogenital tract, Kent, United Kingdom, 1973 | Band-tailed pigeon (*Patagioenas fasciata*), California, 2008 | Human adult female subgingival space, 1959 | Wonga pigeon (*Leucosarcia picata*), Batemans Bay, NSW Australia, 2011 | Barred-shouldered dove (*Geopelia humeralis*), Fingal Head, NSW Australia, 2011 | Domestic rock dove (*Columba livia*), Carabost, NSW Australia, 2011 | Human intestine, Korea, 1950 |
| **Host** | Human | Bird | Human, cat, dog, (bird) | Bird | Bird | Bird | Human, cat, dog, monkey, guinea pig |
| **Platform** | PacBio + HiC | PacBio + HiC | Illumina | Illumina | Illumina | Illumina | Illumina |
| **Estimated genome size (Mb)<sup>a</sup>** | 184.2<sup>b</sup> | 73.0<sup>c</sup> | 104.9 | 77.8 | 81.0 | 68.9 | 100.2 |
| **Assembly size (Mb)<sup>a</sup>** | 181.5<sup>b</sup> | 72.3<sup>c</sup> | 70.4 | 55.4 | 72.5 | 55.7 | 54.3 |
| **Assembly statistics** | six scaffolds<br>(176.6 Mb),<br>212 contigs<br>(4.7 Mb) | six scaffolds | 35,468 contigs | 13,785 contigs | 13,690 contigs | 8,409 contigs | 18,431 contigs |
| **N50 (bp)** | 34.7 Mb<br>(9,240<sup>b</sup>) | 14.5 Mb<br>(11,033<sup>c</sup>) | 11,499 | 18,842 | 22,721 | 28,086 | 10,851 |
| **Repeat content (%)<sup>a</sup>** | 68.6<sup>b</sup> | 32.6<sup>c</sup> | 50.6 | 36.6 | 21.4 | 25.8 | 53.7 |
| **% GC** | 32.7 | 31.2 | 34.1 | 29.8 | 34.4 | 33.7 | 37.8 |
| **No. predicted genes (excluding TEs)** | 37,794 | 28,579 | 29,838 | 23,689 | 33,504 | 24,752 | 26,270 |
| **No. predicted TEs** | 20,720<sup>d</sup> | 625<sup>e</sup> | 459<sup>f</sup> | 254<sup>f</sup> | 150<sup>f</sup> | 164<sup>f</sup> | 897<sup>f</sup> |

<sup>a</sup>Estimated from short-read Illumina sequence data using GenomeScope<sup>70</sup>
<sup>b</sup>Calculated from short-read Illumina sequence data of *T. vaginalis* strain CDC085 (GenBank SRA No. SRX1017343) to provide comparable statistics to other species sequenced using the same sequencing platform
<sup>c</sup>Calculated from short-read Illumina sequence data of *T. stableri* strain CA015840 (GenBank SRA No. SRR30350957) to provide comparable statistics to other species sequenced using the same sequencing platform
<sup>d</sup>Derived from BLAST searches; a TE was counted if it contained at least one gene, which excluded numerous short TE-derived sequences.
<sup>e</sup>Derived from BLAST searches and RepeatModeler.
<sup>f</sup>Estimated using DeviateTE<sup>74</sup>.

Transposable element (TE) sequences, difficult to identify and not classified well in the previous assembly, were meticulously annotated and found to dominate the new *T. vaginalis* reference genome, making up at least 46% of its length (**Table 1, Supplementary** Figure 1). While MULE TEs dominate by total number (7,322), the >4,700 Maverick^11^ (TvMav) TEs comprise >80% of the total TE length and ∼40% of genome length (**Table 2**). We undertook extensive manual curation of TvMavs since they can contain as many as 19 TE genes, lack terminal inverted repeats (TIR), are concatenated or nested within each other, or envelop other types of TEs^11^. Based on several characteristics (length, TIR sequence, gene repertoire, and gene order) of 2,788 well-defined TvMavs, we identified at least three classes: Class 1 (n=902) and Class 2 (n=181) range from 20-25 Kb and differ mainly in TIR sequence, while the abundant and previously undescribed Class 3 (n=1,705) has a bimodal length distribution, suggesting two subclasses with peaks at 10-20 Kb and 23-26 Kb (**Supplementary** Figure 2), as well as a distinct gene repertoire and order (**Supplementary Tables 3-5**).

**Table 2.**
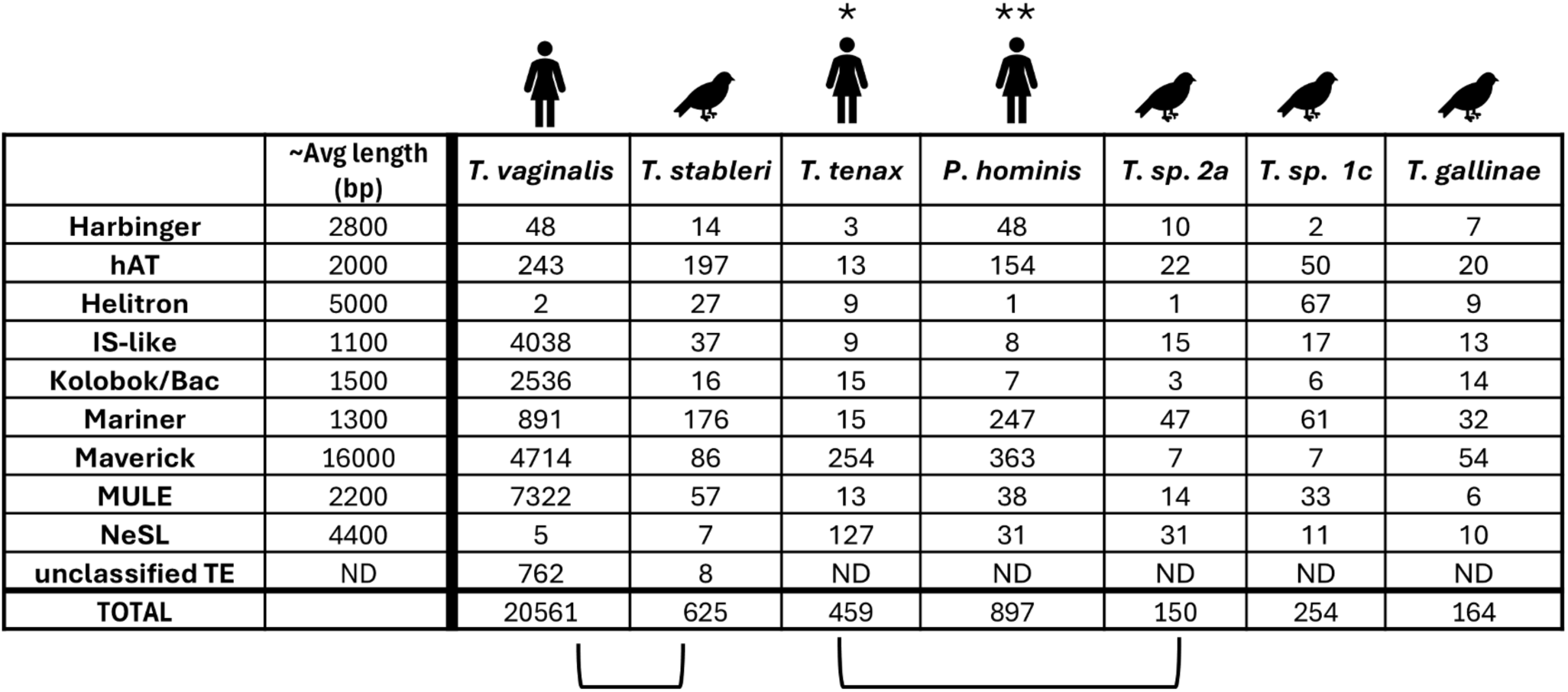
Total number of elements in nine of the most common TE families found in seven trichomonad genomes. Average lengths are derived from those exhibited by *T. vaginalis* TEs. *T. vaginalis* and *T. stableri* genomes were generated by long-read sequencing; all others are short-read assemblies. Icons show host species and bracket endpoints indicate inferred sister species on the phylogenetic tree shown in Supplementary Figure 5. ND: not determined. *\*T. tenax* has been detected in other mammals and in birds. *\*\*P. hominis* has been detected in other mammals.

|  |  |  |  |  |  |  |  |  |
| --- | --- | --- | --- | --- | --- | --- | --- | --- |
|  | ~Avg length (bp) | *T. vaginalis* | *T. stableri* | *T. tenax* | *P. hominis* | *T. sp. 2a* | *T. sp. 1c* | *T. gallinae* |
| Harbinger | 2800 | 48 | 14 | 3 | 48 | 10 | 2 | 7 |
| hAT | 2000 | 243 | 197 | 13 | 154 | 22 | 50 | 20 |
| Helitron | 5000 | 2 | 27 | 9 | 1 | 1 | 67 | 9 |
| IS-like | 1100 | 4038 | 37 | 9 | 8 | 15 | 17 | 13 |
| Kolobok/Bac | 1500 | 2536 | 16 | 15 | 7 | 3 | 6 | 14 |
| Mariner | 1300 | 891 | 176 | 15 | 247 | 47 | 61 | 32 |
| Maverick | 16000 | 4714 | 86 | 254 | 363 | 7 | 7 | 54 |
| MULE | 2200 | 7322 | 57 | 13 | 38 | 14 | 33 | 6 |
| NeSL | 4400 | 5 | 7 | 127 | 31 | 31 | 11 | 10 |
| unclassified TE | ND | 762 | 8 | ND | ND | ND | ND | ND |
| TOTAL |  | 20561 | 625 | 459 | 897 | 150 | 254 | 164 |

### Comparative genomics of seven trichomonads that infect humans, birds, and mammals

We chose several species within genus *Trichomonas* known to be closely related to *T. vaginalis* on the basis of single copy gene phylogenies^12^ for comparative evolutionary studies, including a more distantly related trichomonad species as an evolutionary outgroup. Growing parasites *in vitro* proved challenging and several could not be grown continuously or in sufficient volume to generate the required quantity or quality of DNA for long read sequencing (data not shown). The final list of assembled species and their sequencing statistics is shown in **Table 1**: (1) the New World clade bird parasite *Trichomonas stableri* strain BTPI-3, the closest known relative of *T. vaginalis* sequenced using PacBio/HiC; (2) an Australasian bird parasite *Trichomonas* species genotype 1c (*Trich*. sp. 1c)^8^; (3) the Old World human/mammal/bird parasite *Trichomonas tenax* Hs-4:NIH, (4) an Old World bird parasite *Trichomonas* species genotype 2a (*Trich*. sp. 2a), the closest known relative of *T. tenax*^8^, (5) the Old World bird parasite *Trichomonas gallinae* (TGAL)^8^; and (6) the human/mammal parasite *Pentatrichomonas hominis* (Hs-3:NIH), used as an outgroup for our analyses. Genome size estimates calculated from short reads of the species ranged from 68.9 Mb for *T. gallinae* to 184.2 Mb for *T. vaginalis* (**Table 1**), the latter by far the largest genome size of the seven trichomonad species sequenced. Estimated genome size is associated with host type, being larger in human-infecting species than bird-infecting species, and exhibits a linear relationship (**Supplementary** Figure 3) to estimated repeat content (multicopy genes, TEs, and unclassified repeats; **Figure 4**), which ranges from 21.4% (*Trich.* sp. 2a) to 68.6% (*T. vaginalis*): repeat estimates in avian-infecting species (21%-37%) are far lower than in human-infecting species (51%-69%). Counts of predicted protein-coding genes in the assemblies ranged from 23,689 (*Trich.* sp. 1c) to 37,794 (*T. vaginalis*) and did not display associations with genome size or host type (**Table 1**).

Pairwise whole genome DNA alignments of the species confirmed several previously proposed relationships^6, 7, 8^, including the presence of two lineages that exhibit close ‘sister species’ relationships between a human-infecting species and a bird-infecting species (human *T. vaginalis* with bird *T. stableri*; and human *T. tenax* with bird *Trich. sp.* genotype 2a) (**Supplementary** Figure 4). Whole chromosome synteny mapping of *T. vaginalis* with its avian sister species *T. stableri* showed large differences in chromosome sizes and massive genome rearrangements (**Figure 2**).

**Figure 2.**
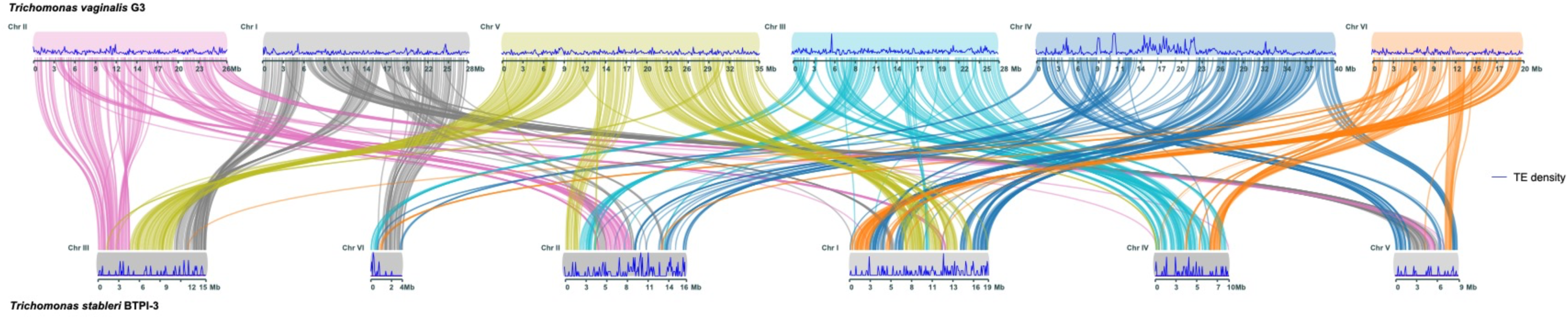
Synteny plot of human parasite *T. vaginalis* and its closest relative in birds *T. stableri.* Each *T. vaginalis* chromosome is colored uniquely, and synteny blocks are indicated by ribbons connecting the species’ chromosomes. TE density (genomic sequence classified as containing TEs) is indicated by normalized density plots in 100 Kb windows at the top and bottom of the figure (blue plots). Chromosomes were reordered for visualization purposes.

We identified 24,465 orthogroups (groups of evolutionarily related genes) across the seven trichomonad species, with 93.8% of all genes being assigned to an orthogroup. Of these orthogroups, 10,457 contain genes from all species (**Figure 3A**), 6,226 contain only single-copy genes, and 2,798 orthogroups, comprising 6.6% of all genes, are species-specific. As expected, the outgroup *P. hominis* contained the largest number of species-specific orthogroups (1,078), followed by *T. vaginalis* (425). We used the 6,226 single-copy orthologs to infer a phylogenetic species tree (**Supplementary** Figure 5).

**Figure 3.**
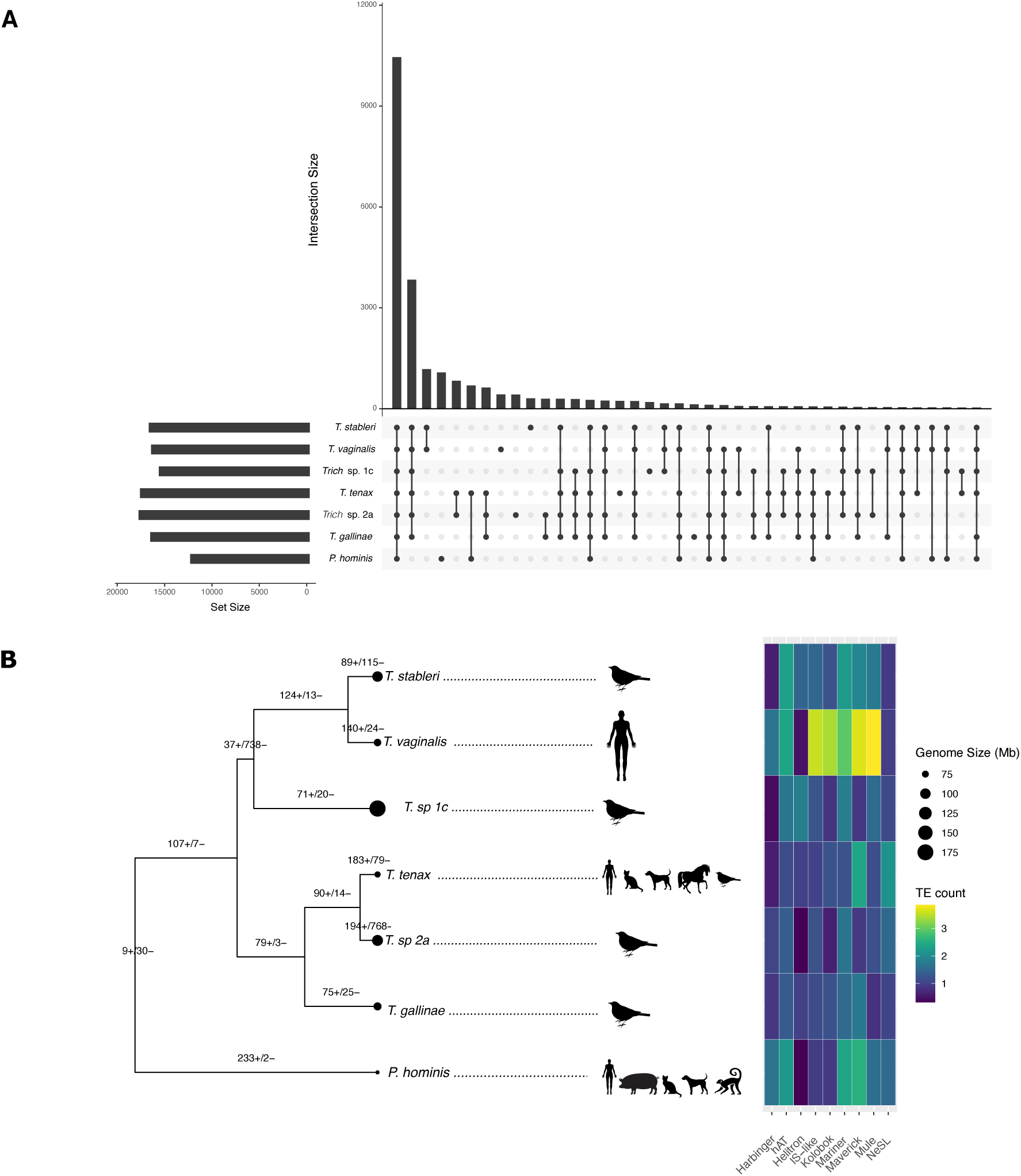
Genome content distribution. **A.** An Upset plot displaying the intersection of orthogroups identified in OrthoFinder across seven trichomonad species. Each vertical bar represents the number of orthogroups shared at each species intersection, the set size indicates the number of orthogroups found in each species, and the connected dots represent the species in the intersection. **B.** Ultrametric tree from 6,226 concatenated single-copy genes. Black dots at terminal nodes are proportional to estimated genome size, and hosts are denoted by cartoons. Estimated gene family expansions / contractions from 12,345 genes are denoted as + or – values on the tree. The heat map shows the log transformed count of TE family members for each tree branch.

The tree strongly supports separate clades for *T. vaginalis* and *T. tenax*, in accordance with the proposal of at least two bird-to-human host switches in the evolutionary history of genus *Trichomonas*^8^. It also resolves the formerly ambiguous placement, from single-gene trees, of Australasian bird parasite *Trich.* sp. 1c among the Old or New World clades^8^; we find strong support for placing *Trich*. sp. 1c with the New World clade.

### The burden of repeats/TEs in trichomonads differs by host type

The correlations between genome size, host type, and repeat content noted above are not necessarily reflected in phylogenetic proximity; for example, the human-infecting trichomonad lineages (*T. vaginalis, T. tenax, P. hominis*), while not closely related by evolution, all appear to have undergone recent and convergent large genome size expansions compared to their avian sister species (**Figure 3B)**. Kmer-based estimates of genome size and repeat content from sequencing reads clearly mark the three human-infecting species as having larger and more repetitive genomes than the bird-infecting species (**Table 1** and **Figure 4**). This is borne out in a comparison of the two long-read assemblies, whose lengths concur with kmer-based estimates, and whose counts of repeat sequences are the most reliable: the major contributor to the much larger genome size of *T. vaginalis* versus *T. stableri* is increased repeat content, particularly expansion of TEs (**Supplementary** Figure 1).

We identified and classified 22,449 TEs in *T. vaginalis*, 3,443 in *T. stableri,* and, within the limits imposed by short-read assembly, 897 in *P. hominis,* 459 in *T. tenax,* and <300 in each of the bird-infecting species (**Table 1**, **Table 2**), again showing a human/bird host disparity. The great majority of TEs identified in all species are Class II DNA transposons, with a single Class I NeSL retrotransposon family identified as particularly abundant in *T. tenax* (**Figure 3B**). Mavericks appear to be more abundant in the three human-infecting species *T. vaginalis, T. tenax,* and *P. hominis* than the bird species, since their size often makes them the dominant TE class by length, even when it is not the most abundant class (**Supplementary** Figure 6), but no pattern of abundance was seen in other TE classes.

A closer inspection of the synteny between *T. vaginalis* with its sister species *T. stableri* in birds (**Figure 2**) revealed the syntenic regions in *T. vaginalis* to be made up of almost equal numbers of TEs (47.3%) and non-TE (52.7%) protein-coding genes, whereas in *T. stableri* the regions are made up of 90.22% non-TE protein-coding genes (**Supplementary Table 6**). Analysis of the *T. vaginalis* protein-coding genes that are not TEs in the syntenic regions revealed many of them to be members of multi-copy gene families enriched in gene ontology (GO) functions such as protein kinases, ATP/GTP binding, and protein phosphorylation. Copy number of several gene families is markedly higher in *T. vaginalis* than *T. stableri*, e.g., BspA-like (73% higher), Saposin-like (SAPLIP) (65%), and leishmanolysin-like proteinase (64%) families (**Supplementary Table 7**), signifying that these expansions were favored in the human host. Most of the remaining gene families, e.g., membrane trafficking proteins, serine peptidase, protein kinases, vary <10% in copy number between the two species, suggesting their gene duplications largely predate the bird-human host switch.

### Evidence of relaxed selection supports a neutral model for genome expansion in human-infecting trichomonad species

To assess levels of genetic drift (a nonadaptive possible driver of expansion of repetitive DNA when selection is relaxed) we used the hypothesis-testing framework RELAX^18^, which asks whether the strength of natural selection has been relaxed or intensified along specified test branches compared to reference branches in a phylogenetic tree. We used the 6,226 single-copy orthologs occurring in all seven species as a proxy for genome-wide sampling of drift. With human-infecting branches as test (foreground) and avian branches as reference (background), we determined which genes evinced significant (p<=0.05) relaxed or intensified selection and found that human-infective branches have more genes under relaxed than intensified selection (n=894 vs. n=494) (**Figure 5A, Supplementary Data file 1**), the converse of the bird-infecting branches.

**Figure 4.**
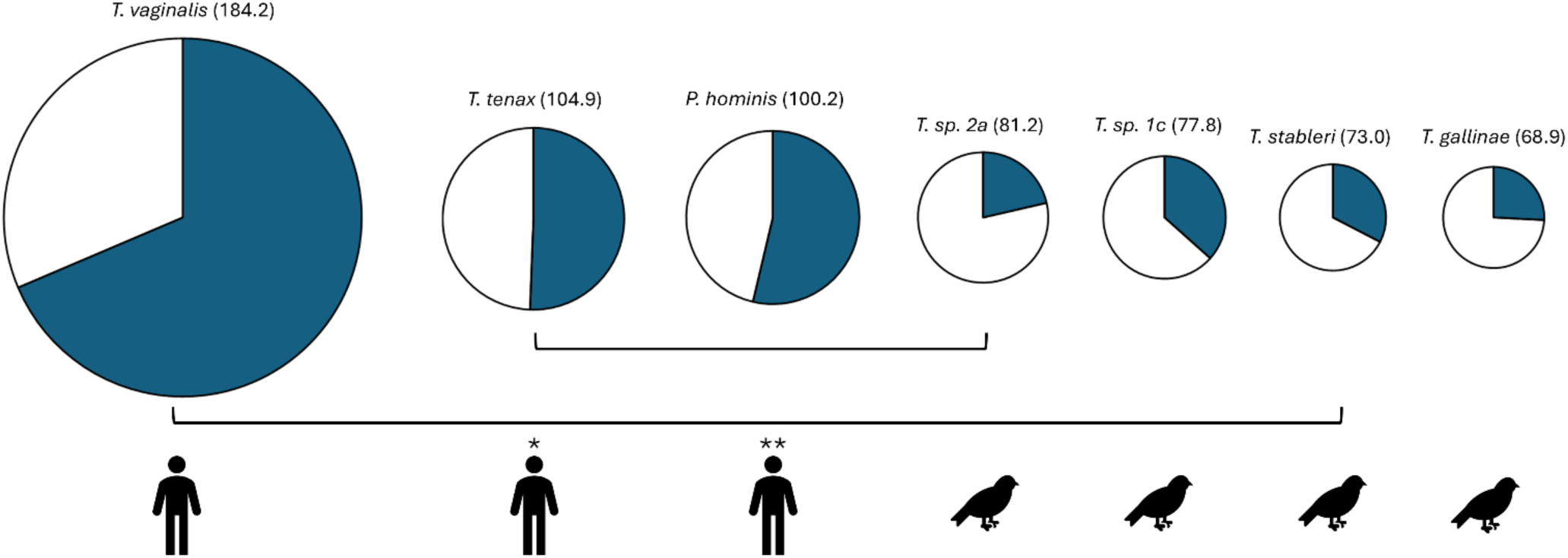
Estimated repeat content length (blue) per estimated genome size. Genomes are shown scaled by their estimated size (Mb, in parentheses). Bracket endpoints denote inferred sister species on the phylogenetic gene tree. Icons indicate human or bird host. *\*T. tenax* has been detected in other mammals and in birds. *\*\*P. hominis* has been detected in other mammals.

**Figure 5.**
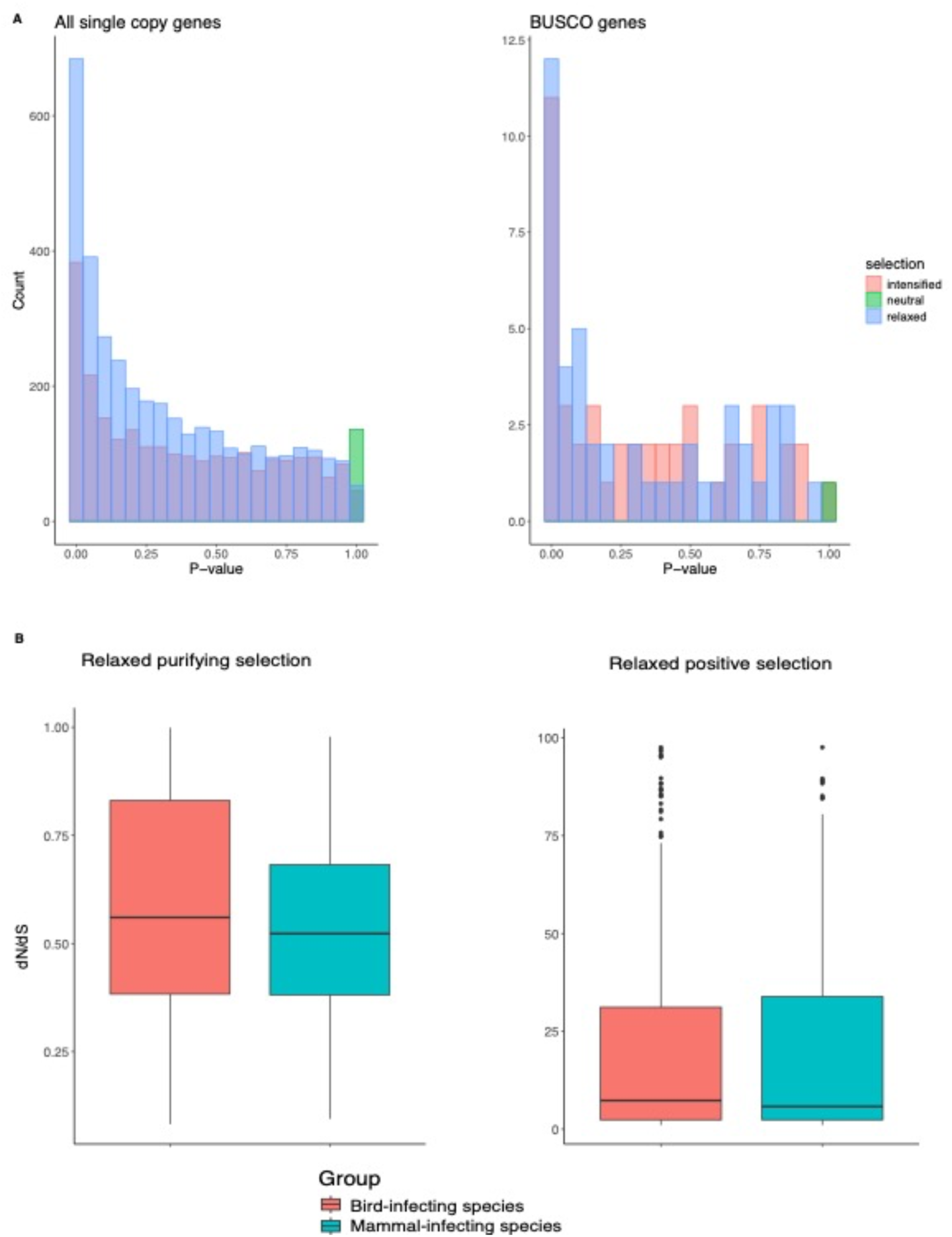
Analysis of orthologs across seven *Trichomonas* species. **A.** Graphs showing count of all single-copy orthologs (SCOs; left panel) and BUSCO genes (right panel) identified by RELAX as being under relaxed, neutral, or intensified selection in the species with expanded genomes (*T. vaginalis* and *T. tenax*) for a range of P-values. **B.** Mean dN/dS values (plotted from 1.0 to 10.0) for SCOs inferred as under relaxed positive selection (left plot) and dN/dS values (plotted from 0.0 to 1.0) for SCOs inferred as under relaxed purifying selection (right plot) for avian-infecting species and mammalian-infecting species.

A gene under relaxed selection may result from an organism switching environments if the gene is obsolete in the new host or tissue, or from increased genome-wide genetic drift (due to changes in parameters such as population size and mode of reproduction)^19^. To rule out host environment as the driver of observed relaxed selection, we used RELAX to test the strength of selection acting on 506 genes from the seven genomes with homology to BUSCO^20^ housekeeping genes, since the rates of evolution of important housekeeping genes are expected to remain constant even in different environments. We found that there are more housekeeping genes under relaxed selection (n=47) than intensified selection (n=44) in the human-infecting species relative to bird-infecting species (**Figure 5A**).

This suggests a role for increased genome-wide genetic drift, rather than relaxed selection targeting genes that are superfluous in the new environment. Consistent with relaxed positive selection, the distribution of average dN/dS ratios (a measure of the strength and mode of natural selection acting on protein-coding genes) for the single-copy orthologs shows a higher median in avian-infecting parasites (7.318) and lower median in human-infecting parasites (5.786) for relaxed purifying selection, and a higher median in avian-infecting parasites (0.585) and lower median in human-infecting parasites (0.542) for relaxed purifying selection (**Figure 5B**). In general therefore, dN/dS in human-infecting trichomonad species has contracted towards 1, i.e., neutral evolution.

### *T. vaginalis* has the largest net gain in number of expanded multicopy gene families compared to other trichomonads

We previously proposed that copy number expansions in *T. vaginalis* multigene families may account for a significant proportion of its unexpectedly large genome size compared to other parasites^9^. We investigated this further by analyzing *T. vaginalis* gene families in the context of our other assembled trichomonad genomes. We used CAFE5^21^, which implements a birth-death model for evolutionary inferences about gene family evolution, to identify multicopy gene families that have expanded or contracted significantly across our trichomonad phylogeny. Of the 26,244 orthogroups, 12,345 (see **Methods**) were analyzed for significant expansions or contractions, of which 3,853 showed significant expansions or contractions in at least one extant species or inferred ancestor (**Figure 3B, Supplementary Data file 1**). We found that among the trichomonad species examined, *T. vaginalis* had the largest net gain (n=116) in number of expanded gene families, consistent with it having undergone the largest genome size increase. These 140 expanded *T. vaginalis* gene families are functionally enriched in gene ontology (GO) terms for transmembrane transport (e.g., ABC transporters), metabolism and translation (**Figure 6)**. We also identified many expanded gene families (n=61) in *T. vaginalis* that have published functions related to parasite pathology, such as host cell adherence^22, 23, 24, 25^, phagocytosis^26^, and extracellular vesicles^27, 28^. We did not find functional enrichment in the 24 multicopy gene families in *T. vaginalis* that have significantly contracted; however, we identified three genes described in previous studies as involved in host cell adherence^22^, phagocytosis^26^, and extracellular vesicles^27, 29^.

**Figure 6.**
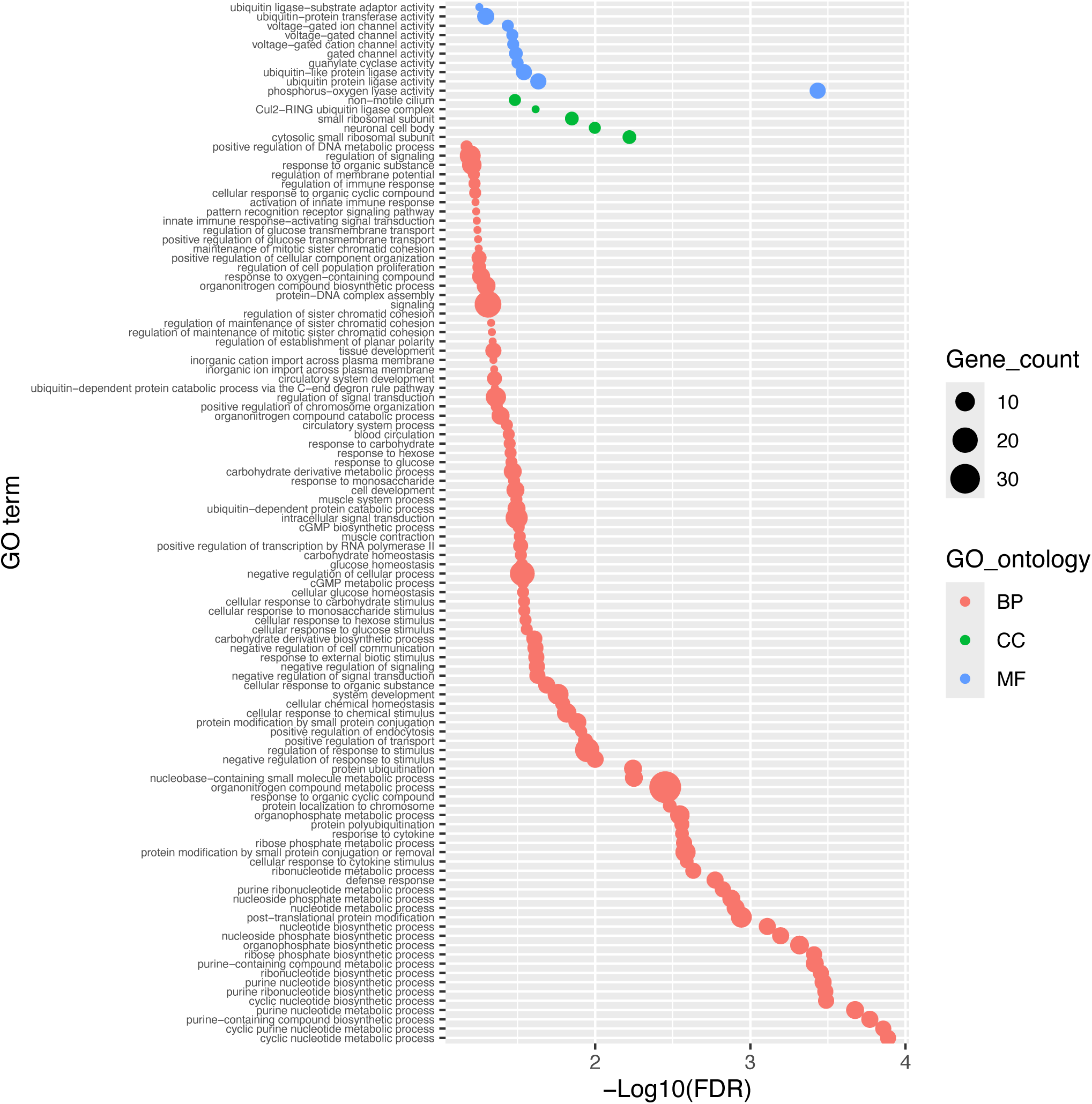
GO enrichment of 140 expanded gene families in *T. vaginalis*. Dot size represents the number of genes with a specific GO term. Biological process (BP), cellular component (CC) and molecular function (MF) are plotted. Only significant GO enrichments after FDR correction (0.05<) are reported.

We found that *T. vaginalis* shares the largest number of expanded gene families not with its bird-infecting sister species *T. stableri*, but with human-infecting *T. tenax* (n=33) and its sister species, bird-infecting *T. sp.* 2a (n=35) (**Figure 7**). These convergently expanded multicopy gene families of *T. tenax* and *T. vaginalis* are enriched for GO terms in metabolism, and include 25 genes previously reported to be associated with cell adherence^22, 23, 30^, microvesicles^27, 29^, and putative virulence factors ^31^. *T. vaginalis* shares the largest number of expanded gene families with *Trich spp.* 2a and consists of GO terms enriched in many biological processes such as transport, telomere maintenance, signal transduction, morphogenesis, and immune response. We similarly found genes with published associations with adherence and microvesicles, but to a lesser degree (n=13). We also identified 15 convergently contracted multicopy gene families in *T. tenax* and *T. vaginalis* species (**Supplementary** Figure 7), but without any enrichment of GO terms.

**Figure 7.**
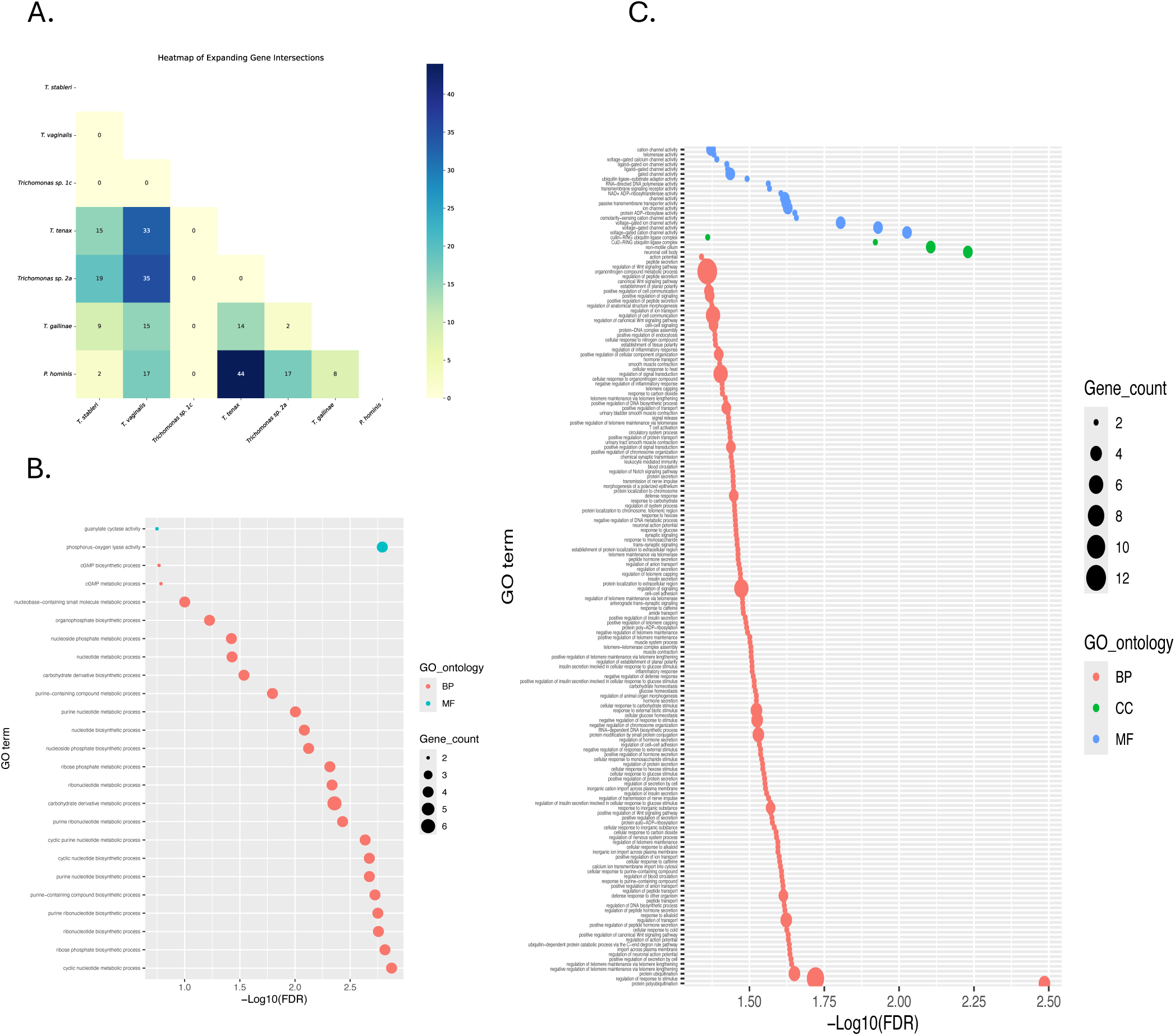
A. Heatmap showing number of shared expanded orthogroups between seven trichomonad species. *T. vaginalis* shares its largest number of expanded orthogroups with *T. tenax* (n=33) and *Trich sp. 2a* (n=35), two non-sister species that infect humans and birds, respectively. **B.** GO enrichment of convergently expanded multicopy gene families in *T. vaginalis* and *T. tenax*. **C.** GO enrichment of convergently expanded multicopy gene families in *T. vaginalis* and *T. sp. 2a*. For both **B** and **C**: size of dot represents the number of genes with a specific GO term; biological process (BP), cellular component (CC) and molecular function (MF) are plotted; only significant GO enrichments after FDR correction (0.05<) are reported.

### Evolution modeling identifies *Trichomonas* genes under positive selection and genes putatively involved in the bird-to-human host switch

We used a branch-site model implemented in aBRASEL^32^ to test for positive selection in single-copy genes of our trichomonad species (**Supplementary** Figure 8**, Supplementary Data file 1**).

Approximately 27% of genes with evidence of being under positive selection were shared between two or more species, the rest being specific to a single species. The shared genes were enriched in GO terms for translation, intracellular transport, and cytoskeleton/motility, most likely reflecting functions essential to trichomonads generally (**Supplementary** Figure 9). A relatively large number of these shared genes have been previously associated with phagocytosis^26^ (n=44) and include proteases, cytoskeleton genes, transmembrane and transporter genes, vesicular trafficking, and metabolism-related genes, and a similar number were associated with microvesicles^27^ (n=40), including a number of tRNA synthetases and peptidases, regulatory and binding proteins. Smaller numbers of shared genes were associated with adherence^22, 24, 30, 33^ (n=21) and included transporters and membrane proteins; exosomes^29^ (n=9) including one core exosomal protein; and proteins of the secretome^28^ (n=4), and carbohydrate-active enzymes (CAZymes, n=1) implicated as virulence factors^31^.

We identified 138 candidate genes with evidence of positive selection in *T. vaginalis*, 69 of which are unique to the *T. vaginalis* lineage. While no GO terms were found to be enriched among them, ten genes (TVAGG3_0302500, TVAGG3_1001150, TVAGG3_1088290, TVAG_005750, TVAG_062520, TVAG_117090, TVAG_152520, TVAG_313880, TVAG_437950, TVAG_453350) **Supplementary Data file 1**) are specific to *T. vaginalis* and have experimentally verified functions associated with adherence^22, 23, 25^, microvesicles^27^, the secretome^28^, phagocytosis^26^, and CAZymes^31^. *T. tenax* shows 45 genes with evidence of positive selection, 26 of which are unique to the lineage. We did not find GO enrichment among these 45 genes. However, six genes (TVAG_097660, TVAG_127300, TVAG_137880, TVAG_237760, TVAG_270770, TVAG_459530) under positive selection and shared between other trichomonad species have been associated with adherence^24^, microvesicles^27^, exosomes^29^, and phagocytosis^26^; all of these genes are shared with *T. vaginalis*.

Assuming that trichomonads have independently host-switched twice from birds to humans to generate the *T. vaginalis* and *T. tenax* lineages^8^, and that selection will act on similar genes when different lineages independently adapt to similar environments, we applied the convergent evolution model RERconverge^34^ to identify single-copy genes putatively involved in the transition to a human host (**Supplementary Data file 1**). Convergent evolutionary rate shifts can indicate whether changes in selection in a gene cohort are due to deaccelerated evolution (i.e., purifying selection) versus accelerated evolution (i.e., relaxed or positive selection) compared to the average rate across the phylogeny. Of 6,226 single-copy orthologs, 320 showed evidence of convergent deaccelerated evolution in the human-infecting branches *T. vaginalis* and *T. tenax*. Several of these genes are reported to be associated with phenotypes of phagocytosis^26^ and adherence^22, 30^, as well as microvesicle^27^ and exosome^29^ structures, and CAZymes^31^. A total of 93 single-copy orthologs showed evidence of convergent accelerated evolution in *T. vaginalis* and *T. tenax*; several of these have been reported to be involved in adherence^30^, phagocytosis^26^, and microvesicle-like structures^27^. Overall, genes involved in

adherence, phagocytosis, and microvesicle-like structures are implicated in the transition from a bird to a human host.

## Discussion

We present here a comparative analysis of the first chromosome-scale genomes of *T. vaginalis*, an extracellular microbe that causes the most common sexually transmitted parasitic infection of humans, and its sister species in birds, *T. stableri*, with genomes from five other species of human- and bird-infecting trichomonads. These comparisons illuminate for the first time differences in protein-coding gene and TE content, genomic architecture, and gene evolution across the trichomonad phylogeny, and identify genes implicated in the inferred spillover event from avian to human host.

All of the trichomonads we sequenced (indeed all trichomonads and tritrichomonads) have much larger genomes than other orders of single-celled parasites that cause important human diseases^10, 35^. The major contributor to increased genome size is increased repeat content, in particular TE expansion which has been proposed to be triggered by major environmental changes^36^. TEs constitute the bulk of repetitive DNA in *T. vaginalis*, and likely the other genomes presented here as well. The presence of the same classes of TEs in all of the trichomonad genomes points to either multiple invasions of an ancient common ancestor or multiple invasions and expansions after divergence. For example, the very high sequence similarity of the hundreds of Mariners we recently reported in *T. vaginalis* points to their recent expansion in that genome^37^, and high polymorphism in Mariner insertion sites across different *T. vaginalis* strains also suggests recent active transposition of this TE class in the species^37^. At least 45% of the *T. vaginalis* genome length is made up of three classes of long Maverick DNA transposons (TvMavs), an ancient DNA transposon lineage likely derived from plasmids, virophages and giant viruses. Mavericks in nematodes were recently found to be agents of cross-species horizontal transfer of non-TE gene ‘cargo’^38^; we observed non-TE genes occurring sporadically inside TvMavs which could represent cargo genes (data not shown). A greater abundance of TvMavs in the human-infective species *T. tenax*, *T. vaginalis*, and *P. hominis* also appears to be the main contributor to their genome size increases relative to bird-infecting species. A ‘transposome’ analysis across species is needed to clarify the likely complex evolutionary history of trichomonad TEs, and to extend the previous studies on the TE transcriptional silencing mechanisms elucidated in some *T. vaginalis* TE families^39^.

We previously hypothesized that the genome size expansion of *T. vaginalis* reflected a relaxation of selection when the parasite underwent a population size bottleneck during its transition from a GI environment to the urogenital tract^9^. In the present study, we found an overall trend of relaxed selection amongst human-infecting compared to bird-infecting *Trichomonas* species, suggesting higher levels of genetic drift as a factor in their genome expansion. But do host-switch bottlenecks alone account for the relaxation of selection? Peters et al.,^8^ estimated the co-divergence between host and parasite and observed relatively shallow branches in the parasite tree, indicating recent divergence in the parasites but not the hosts. Additionally *T. gallinae*, and *Trichomonas sp. 2a* have been identified across bird orders, not just genera^8^. These observations suggest that recent host shifting, including across fairly large evolutionary distance, is a general phenomenon amongst columbid *Trichomonas*, and that we would therefore expect bottlenecks (and relaxed selection) in these species as well, if the hypothesis is true. But the relative lack of relaxed selection we observed in columbid *Trichomonas* overall suggests that factors other than bottlenecks contributed to the host-associated difference in selection strength.

Mode of parasite reproduction is one such possible contributor. Asexual reproduction can lower the effective population size through decreased genetic variation and global reduction of variation due to background selection and genetic hitchhiking^19^. The last common ancestor to eukaryotes is thought to have reproduced sexually, and among extant eukaryotes sexual reproduction is generally the norm. We previously accumulated evidence that *T. vaginalis* may have undergone sexual recombination in its evolutionary past^40^; a putative hybridization event has also been described in *T. gallinae*^41^, raising the possibility that sex occurs in other *Trichomonas* species. Thus it could be that a shift from sexual to asexual reproduction, in addition to a bottleneck, accompanied host switching, and facilitated relaxed selection, catalyzing large-scale structural changes to the genomes. Further investigation of reproduction in genus *Trichomonas* is needed to confirm this hypothesis.

Among the trichomonads in our study, we found the largest net gain in number of expanded gene families in *T. vaginalis*, and the highest number of gene family expansions shared with *T. vaginalis* in the *T. tenax/T. sp. 2a* clade, indicating that the latter similarity results from convergent evolution. The set of expanded families shared by *T. vaginalis* and human-infecting *T. tenax* is different from that shared by *T. vaginalis* and bird-infecting *T. sp. 2a*. Families that expanded in *T. vaginalis* and *T. tenax* feature more genes involved specifically in metabolism, cell adherence, microvesicles, and virulence, than those expanded in *T. vaginalis* and *T. sp. 2a*, which could be evidence for human- (or at least mammal-) specific adaptations. Indeed, the diverse array of glycoside hydrolases, Carbohydrate Active enZymes (CAZymes), and carbohydrate-binding modules, identified through a recent comparative analysis of *T. vaginalis* and *T. tenax*^31^ are likely shared virulence factors that potentially target host or bacterial glycans, and induce and/or amplify damaging inflammation and bacterial dysbiosis, known to exacerbate periodontitis and vaginitis. The functions associated with the shared *T. vaginalis/T. sp. 2a* families, on the other hand, could be those useful to parasitic trichomonads with bird-host ancestors. The recent reports of *T. tenax* in birds^2^ complicates this hypothesis. However, that report is based upon genotyping of the multicopy ITS1/5.8S/ITS2 rRNA small subunit gene, where discrimination between species can be based on as little as <=1% difference in sequence identity. Moreover, our *T. tenax* genome sequence is of a strain isolated from a human subject and presumably adapted to that host. More sequence data from *T. tenax* isolates from humans and birds are needed to clarify this.

The columbid upper GI tract and the oral and vaginal cavities of humans are lined with stratified, non-cornified epithelia^42, 43^, a histological similarity that conceivably enabled the ancestral colonization of a human tissue by a bird trichomonad. At the same time, convergent changes in the human-infective species suggest there was enough microscale difference in the host environments to drive adaptation.

Convergently evolving multicopy gene families in *T. vaginalis* and *T. tenax* included some associated with cell adherence, suggesting specifically that differences in surface membrane proteins in bird versus human mucosal epithelium could foster selection for differential adherence to host tissues.

Multicopy genes are challenging to use in some evolutionary analyses because it is difficult to identify orthologues between them. But evidence from analysis of single-copy gene evolution can illuminate phenomena such as host-switching or spillover. We looked at single copy orthologues two ways, specifically for positive selection, and more generally for rates of evolution indicating purifying (deaccelerated evolution) versus relaxed or positive selection (accelerated evolution). Most single-copy genes with evidence for positive selection were species-specific, suggesting fine-tuning of the parasite to particular environments. Single-copy orthologues in *T. vaginalis* and *T. tenax* identified as displaying convergent deaccelerated or accelerated evolution were often related to the endo- and cell membrane systems, and also adherence, phagocytosis, and mitosis. The endomembrane system generates extracellular vesicles, e.g., exosomes and microvesicles, which have been shown in *T. vaginalis* to prime host cells for adherence, modulate the host’s immune response, facilitate cell-to-cell communication, and promote host cell colonization^23, 25, 27, 29^. Convergent selection for endomembrane system genes could reflect adaptation of the parasite to the host’s surface membranes and immune system; vesicles can carry cargo that affect host gene expression, and the removal of these vesicles from the extracellular milieu reduces the adherence of the parasite to host cells^23^. Parasite adherence to host mucosal cells is essential in establishing an infection, and parasite phagocytosis is involved in nutrient acquisition^44^ and immune cell evasion^45^. Autophagy is associated with the pathogenicity of several protozoan parasites and has been demonstrated to increase the survivability of *T. vaginalis* under nutrient starvation^46^ as well as participate in proteolysis^47^. Both phagocytosis and autophagy also involve the endomembrane system. Peculiarly among eukaryotes, *T. vaginalis* mitosis can occur during phagocytosis, which has been hypothesized to be selectively advantageous for a parasite in a hostile environment with scarce nutrients^48^. Combined, our results identify and highlight these candidate genes and gene families implicated in the spillover of parasites from the upper GI tract of columbids into the human reproductive tract for further investigation into trichomonad evolution and adaptation to human hosts.

## Methods

### Generation of a *T. vaginalis* chromosome-scale assembly and annotation

DNA was extracted from *T. vaginalis* strain G3 parasites cultured in modified Diamond’s media and sequenced using Pacific Biosciences Inc. sequencing chemistry on 56 SMRT cells using the PacBio RSII instrument, generating 2,043,705,869 reads that were initially assembled using FALCON^49^. The initial assembly had a total span of 173 Mb across 1194 contigs with a contig N50 size of 321 Kb. Hi-C library preparation and sequencing were performed as described^50^, and PBJelly^51^ used to close any scaffold gaps. In total, this yielded six chromosome-scale scaffolds containing 97.4% of the original assembly with a scaffold N50 size of 27.3 Mb, scaffold N90 size of 20.0 Mb, and improved the contig N50 size to 444 Kb. Pilon^52^ was used for assembly polishing two times using published G3 Illumina reads (SRA# SRR4734558), Sanger reads^9^, and RNA-seq reads^39^ mapped to the assembly using BWA^53^.

Structural annotation used BRAKER2^54^, STAR^55^-mapped RNAseq reads, and a training set of 539 high-confidence *T. vaginalis* protein sequences. *De novo* structural annotation was augmented by gene model transfer from the 2007 *T. vaginalis* assembly (TrichDB release 52), using Liftoff^56^ with parameters *-s 0.9* and *-a 0.9*. Functional annotation used one of six criteria: (1) identity to proteins with previously experimentally characterized function; (2) identity to proteins previously inferred as horizontally transferred from firmicute bacterium *Peptoniphilus harei*^17^; (3) strong similarity (90% ID over 90% length) to UniProtKB/Swissprot entries; (4) orthology group membership and function using eggnog-mapper^57^ and the eggNOG database of orthology groups^58^; (5) protein domains returned by Interproscan (v 5.52.86)^59^; (6) DeepFRI function prediction from predicted protein structure^60^; and for the remainder (6) DeepGOPlus^61^ version 1.0.20 function prediction. Proteins that could not be assigned a function by these means were called ‘conserved hypothetical’. GO enrichment analysis was undertaken using the hypergeometric distribution incorporated into an inhouse Python script.

Maverick TEs were identified by BLAST using ORFs from 11 ‘canonical’ Mavericks identified previously^11^. Ordered blocks of Maverick ORFs were marked in the polished assembly. Canonical and novel terminal inverted repeats (TIRs) flanking the blocs were identified with BLAST and Inverted Repeats Finder^62^. Other TE families were identified through BLASTn queries using consensus TE sequences from Repbase^63^, GyDB^64^, and a custom database of previously identified TEs; RepeatModeler2^65^ was used to identify novel potential TEs. We used phylogenetic analysis, motif identification, and Interproscan to validate the classification of RepeatModeler TE consensuses, and einverted (EMBOSS^66^) and GenericRepeatFinder^67^ to annotate TIRs and target site duplications followed by manual inspection of a multiple sequence alignment of the TE family.

### Sequencing, assembly, and annotation of six additional trichomonad species

*T. gallinae* strain TGAL, *Trichomonas* species genotype 1c, *Trichomonas* species genotype 2a, *T. tenax* strain Hs-4:NIH (ATCC 30207), *P. hominis* strain Hs-3:NIH (ATCC 30000), *T. stableri* strains CA015840 (ATCC PRA-430) and BTPI-3 (ATCC PRA-412) were grown axenically *in vitro* under standard conditions, DNA extracted, libraries generated and sequenced on an Illumina HiSeq platform. *T. gallinae* strain TGAL, *Trichomonas* species genotype 1c, *Trichomonas* species genotype 2a, *T. tenax* strain Hs-4:NIH (ATCC 30207), *P. hominis* strain Hs-3:NIH (ATCC 30000), *T. stableri* strain CA015840 (ATCC PRA-430) were grown axenically *in vitro* under standard conditions, DNA extracted, libraries generated and sequenced using paired-end (mean distance=250 bp) and mate-pair (mean distance=5500 bp) sequencing on an Illumina HiSeq platform. Barcode sequences were trimmed using the fastx toolkit (https://github.com/agordon/fastx_toolkit), sequencing errors were corrected using Quake^68^ and then assembled using SOAPdenovo2^69^ yielding contig N50 sizes of 10 Kb to 25 Kb (**Table 1**). Genome sizes were estimated from Illumina reads using GenomeScope^70^. *T. stableri* strains BTPI-3 and CA015840 were sequenced using Pacific Biosciences Sequel II SMRT technology, assembled using hierarchical genome-assembly HGAP (Pacific Biosciences, SMRT Link V11.1) and Canu^71^, and the resulting assemblies scaffolded using Hi-C data^50^. RNA-seq data were generated in triplicate for *T. stableri* strains BTPI-3 and CA015840, using total RNA extracted from three biological replicate cultures for each strain, stranded mRNA preparation, and the resulting libraries run in HighOutput mode on a NextSeq 500 sequencer to produce 2 x 75 bp paired-end reads. TE expression was estimated in *T. stableri* as for *T. vaginalis* G3 above.

*De novo* gene finding and annotation of the remaining five assemblies was performed using AUGUSTUS^72^ with *ab initio* training using the standard translation code. RNA-seq data were used for transcript assembly and annotation where available for each species, and annotation was manually curated when possible. For annotation of TEs, we used BLASTn^73^ with conserved sequence motifs of all TE consensus sequences identified in *T. stableri* and *T. vaginalis* as queries. For the other species, their lower assembly quality precluded annotation of TE sequences. To quantify them, TE queries based on *T*.

*vaginalis/T. stableri* consensus sequences were used in BLASTn to find matches in each species, which were used to generate per-species consensus sequences. Raw reads were mapped to these consensus sequences using deviaTE^74^, to estimate the true insertion frequency of each TE family except Mavericks, where raw reads mapping to the integrase ORF were used to estimate TE frequency.

### Comparative genomics

DNA sequence similarity across the seven *Trichomonas* species at the whole genome level was calculated using MUMmer v.3.23 dnadiff algorithm^75^ using default parameters, and Bray-Curtis dissimilarity statistics. Synteny analysis was determined using MCScanX^76^ using default parameters (Maximum Evalue: 1e-10, Num.of BlastHits: 5 [minimum collinearity length]), which identifies paralogous, orthologous, and single-copy genes using amino acid sequences of annotated ORFs and identifies collinear blocks of genes between species; a synteny/collinear block consists of ≥ five genes conserved between the two species. OrthoFinder^77^ was used to identify 6,226 single-copy orthologs (SCOs) across the seven trichomonad species, and GO terms assigned to them using embedding similarity^78^. Other analyses used custom in-house Python Scripts and packages in R, such as UpSetR in version 1.3.3; 2017.

### Phylogenetic and evolutionary analyses

A species tree of the seven trichomonad species was generated from 6,226 genes present in one copy in each genome (‘single-copy orthologues’). Orthologues were aligned using PRANK^79^ with default parameters and concatenated to generate a supergene matrix for phylogenetic inference with Phangorn^80^, estimating the best evolution model as GTR + G + I using AICc and executing 1000 bootstraps for analysis. To test if expanded genomes experienced genome-wide relaxed selection, we used RELAX^18^ on the 6,226 single-copy orthologs. We tested human-infecting *Trichomonas* species (*T. vaginalis and T. tenax*) with the four avian-infecting species set as background, and the outgroup *P. hominis* excluded. We tested the avian sister species (*T. stableri* and *Trichomonas* sp. genotype 2a) of *T. vaginalis* and *T. tenax* against all *Trichomonas* species (i.e., excluding *P. hominis*). BUSCO^20^ was used to identify single-copy orthologs that are near-universal across eukaryotes. Significant genes in RELAX results were searched against a curated database of published papers associated with specific phenotypes such as virulence. CAFE5^21^ was used to implement a birth-death model for evolutionary inferences about gene family evolution. A total of 12,345 of 26,244 orthogroups were tested that met the CAFÉ requirement that each orthogroup include the outgroup species *P. hominis*. The R package RERConverge^34^ was used to test for association between relative evolutionary rates of genes and the evolution of traits across the phylogeny. This enabled the generation of lists of candidate genes associated with evolutionarily important traits. aBRASEL^32^ was used to test for positive selection in single-copy genes of *Trichomonas* species using default parameters.

### Data availability

Data supporting the findings of this work are available within the paper and its Supplementary Information files. The raw sequence data have been deposited in GenBank under the following accession numbers: *T. vaginalis* strain G3: Bio-Project PRJNA885811, Genome Accession SAMN31107788, RNA-Seq SUB12510628; *T. stableri* strain BTPI-3: Bio-Project PRJNA816543, Genome SUB11194301, RNA-seq SUB12511058; *T. stableri* strain CA015840: Bio-Project PRJNA828130, Genome SUB11350628, RNA-seq SUB12511233, Illumina DNA-seq reads SRA SRR30350957; *T. gallinae* strain TGAL: Bio-Project PRJNA885811, Genome SUB14002162; *Trichomonas* spp. genotype 1c: Bio-Project PRJNA885811, Genome: SUB14002711; *Trichomonas* species genotype 2a: Bio-Project PRJNA885811, Genome SUB14002751; *T. tenax* strain Hs- 4:NIH: Bio-Project PRJNA885811, Genome: SUB14002737; *P. hominis* strain Hs-3:NIH: Bio- Project PRJNA885811, Genome SUB14002786.

## Acknowledgements

We thank Mari Shiratori, Sally D. Warring, Martina Bradic, Akash Sookdeo, Charlotte Darby, and Srividya Ramakrishnan for initial wet lab and genome analyses. We thank Ellen Pritham for supplying canonical Maverick sequences. Research reported in this publication was partially supported by: the NYU IT High Performance Computing resources, services, and staff expertise; CMRPD1M0571-2 from Chang Gung Memorial Hospital and NSTC–110–2320B–182–016–MY3 from National Science and Technology Council, Taiwan; Australian Government Wildlife Exotic Disease Preparedness Program; and the National Institute Of Allergy And Infectious Diseases of the National Institutes of Health under Award Number R21AI149449 and U24AI183870. The content is solely the responsibility of the authors and

does not necessarily represent the official views of the National Institutes of Health.

## Author contributions

J.M.C. and M.C.S. designed the study, provided the funding for resequencing *T. vaginalis*, supervised the work, and interpreted the analyses. Y.A.G., C.K.J., K.H.R. and R.G. provided *Trichomonas* isolates. P-J.J., Y-M.Y., C-C.L, H.L, T-W.C., C-H. C, and P.T. provided the *T. tenax*, *P. hominis, T. stableri* CA015840 Illumina whole genome sequence data, and A.P. and S.R. provided the *T. gallinae*, *Trichomonas* spp. genotype 1c, and *Trichomonas* sp. genotype 2a Illumina whole genome sequence data. S.A.S, J.C.O, F.C.H., F.B., T.R-B., and H.L. undertook genome annotation and analysis, and additionally F.B. and F.C.-H. generated the *T. stableri* BTPI-3 genome sequence and undertook wet lab work. C.C., D.B., V.G., and R.A.B. helped with *T. vaginalis* functional annotation. All authors contributed to the writing of the manuscript and approved the final version before submission.

## Additional Information

**Supplementary information** The online version contains supplementary material available at …

**Correspondence and requests for materials** should be addressed to Jane M. Carlton

